# Pre-movement respiration–action coupling is specific to self-initiated action: Evidence for action alignment with ongoing breathing

**DOI:** 10.64898/2026.02.05.704105

**Authors:** Hiroshi Shibata, Hideki Ohira

## Abstract

Voluntary actions are coupled with the respiratory cycle, yet whether this coupling reflects action timing aligning with breathing or breathing adjusting to anticipated action remains unclear. This pre-registered study compared respiration–action coupling across three conditions using a modified Libet clock task. Thirty participants performed key presses under Self-initiated (freely chosen timing), Delayed (predictable timing after one full clock rotation), and Immediate (speeded response to stimulus onset) conditions. Circular uniformity tests revealed significant respiratory phase concentration at key press exclusively in the Self-initiated condition, and time-resolved phase consistency analysis showed that coupling emerged before movement onset only in this condition. These findings suggest that action timing aligns with ongoing breathing rhythms, and that temporal freedom, rather than preparation time or predictability, is critical for this coupling to emerge. Additionally, actions performed during inhalation were followed by shorter and deeper subsequent breaths across all conditions, indicating greater perturbation of ongoing breathing when acting during inhalation. Participants reported a subjective preference for pressing during exhalation, which correlated with individual differences in exhalation bias during self-initiated actions. Body awareness, but not interoceptive awareness or trait anxiety, predicted coupling strength. These results suggest that pre-movement respiration–action coupling reflects an implicit process whereby self-chosen action timing exploits ongoing respiratory states, potentially minimizing action-related respiratory perturbation.

## 1 Introduction

Breathing is a bodily rhythm that influences perception and behavior. Accumulating evidence indicates that respiratory phases—particularly inhalation and exhalation—modulate perception, cognition, and emotional processing (Mizuhara & Nittono, 2023; Tort et al., 2018; Zelano et al., 2016). The neural basis for these effects has been increasingly clarified: respiratory rhythms entrain brain-wide neural activity (Kluger & Gross, 2021; Tort et al., 2025), and cortical excitability in both visual (Kluger et al., 2021) and motor cortex (Engelen et al., 2025) fluctuates with the respiratory cycle.

Breathing couples with external events and voluntary actions. Respiratory cycles tend to be synchronized with the structure of cognitive tasks and the predictable onset of sensory stimuli (Harting et al., 2025; Johannknecht & Kayser, 2022; Saltafossi et al., 2025). This synchronization has been observed across diverse sensory modalities, including visual (Chalas et al., 2025), auditory (Goheen et al., 2024), and tactile domains (Grund et al., 2022). Moreover, when individuals freely initiate cognitive tasks, they tend to do so at specific respiratory phases, suggesting that respiratory timing is linked to the initiation of self-paced behavior (Perl et al., 2019). Respiratory coupling has also been investigated in the context of voluntary action itself. Early work demonstrated that spontaneous finger movements tend to occur at specific respiratory phases (Rassler, 2000). Park et al. (2020) further investigated this coupling using the Libet clock task (Libet et al., 1983), in which participants press a key at a self-chosen moment while watching a rotating dot, and found that key presses were biased toward the exhalation phase and associated with the readiness potential. Subsequent research extended these findings to cognitive tasks such as motor and visual imagery, demonstrating similar respiratory coupling patterns (Park et al., 2022). Shibata and Ohira (2025) showed that respiration–action coupling generalizes across different movement types and effectors, with a consistent exhalation bias.

However, respiration–action coupling in voluntary actions raises several unresolved questions. First, the directionality of this coupling remains unclear: does action timing align with ongoing breathing rhythms, or do individuals adjust their breathing in anticipation of action? In self-paced paradigms, these two possibilities are difficult to disentangle because both processes could produce similar phase-locking patterns (Rassler, 2000; Rassler & Raabe, 2003). If breathing adjusts to anticipated action, such adjustment should also occur when action timing is predictable and sufficient preparation time is available, even without temporal freedom. Conversely, if coupling requires the freedom to choose when to act (Galvez-Pol et al., 2025; Lopez-Martin et al., 2026), it should emerge only when individuals can select action timing based on their ongoing physiological states. Second, although exhalation bias has been consistently reported (Park et al., 2020, 2022; Shibata & Ohira, 2025), its functional significance remains poorly understood. Third, substantial individual differences in coupling strength and direction have been reported (Shibata & Ohira, 2025), yet what factors contribute to this variability remains to be clarified.

The present study addressed these questions by comparing respiration–action coupling across three conditions using a modified Libet clock task. In the Self-initiated condition, participants freely chose when to press a key, allowing both temporal freedom and preparation time. In the Delayed condition, participants pressed the key at a predictable, externally determined moment (one full rotation of a clock dot), providing preparation time and predictability without temporal freedom. In the Immediate condition, participants pressed the key as quickly as possible upon stimulus onset, providing neither preparation time nor temporal freedom. Because stimulus onset occurred at random respiratory phases in both the Delayed and Immediate conditions, any respiratory phase concentration at key press in these conditions would reflect respiratory adjustment rather than action selection based on ongoing phase. If breathing adjusts to anticipated action, coupling should emerge in both the Self-initiated and Delayed conditions. If, however, action timing aligns with ongoing breathing rhythms, coupling should emerge only in the Self-initiated condition, where temporal freedom permits action selection based on respiratory phase. Additionally, we explored the functional significance of the exhalation bias by examining respiratory changes following actions performed during different respiratory phases. We reasoned that post-action respiratory changes may serve as a proxy for the physiological cost of acting at different phases. We also examined the relationship between implicit coupling and subjective action preference, and investigated whether individual differences in interoceptive awareness, body awareness, and trait anxiety predict coupling strength.

## 2 Methods

### 2.1 Participants

30 healthy Japanese university students (22 female; mean age = 20.77, *SD* = 2.27 years) from Nagoya University participated in this study. One participant was excluded due to technical problems during data acquisition, resulting in a final sample of 29 participants. An additional participant was excluded from the Delayed condition due to insufficient valid trials (see Section 3.1 for details), resulting in 28 participants for analyses involving comparisons across all three conditions. All participants were right-handed. None reported a history of neurological, psychiatric, or respiratory disorders. Sample size was determined based on previous studies examining respiratory-brain coupling during voluntary movements, which typically included 20-30 participants (Park et al., 2020; Shibata & Ohira, 2025). We recruited 30 participants to account for potential dropouts. The study was approved by the Ethics Committee of the Department of Psychology at Nagoya University (No. NUPSY-250728-G-01) and was conducted in accordance with the Declaration of Helsinki. All participants provided written informed consent prior to participation and received monetary compensation. This study was pre-registered on AsPredicted (#240669). Minor deviations from the preregistration are documented in the Supplementary Materials (S1).

### 2.2 Physiological recording

Respiration data were recorded continuously using an MP160 data acquisition system with an RSP100C respiratory amplifier (Biopac Systems Inc.) at a sampling rate of 200 Hz. Respiratory movements were measured using a respiration belt transducer (TSD201; Biopac Systems Inc.) placed around each participant’s abdomen. Electrocardiogram (ECG) data were also recorded using the MP160 system with an ECG100C amplifier (Biopac Systems Inc.) at a sampling rate of 200 Hz. Disposable electrodes were placed below each clavicle and on the lower left abdomen. All physiological data were acquired using AcqKnowledge-NDT software (Biopac Systems Inc.). Participants were instructed to breathe naturally through the nose throughout the experiment.

### 2.3 Experimental tasks

#### 2.3.1 Libet clock task

Participants performed a modified Libet clock task under three conditions: Immediate (reactive), Delayed (predictable), and Self-initiated (spontaneous). In all conditions, a clock face was displayed on the screen with a dot rotating around the clock at a constant speed of 5,120 ms per cycle. Participants pressed the key with their right index finger in all conditions.

##### Immediate condition (reactive key press)

Each trial commenced when a black dot appeared at a random position on the clock face and began rotating. Participants were instructed to press the key as quickly as possible upon the dot’s appearance. The rotating dot disappeared immediately upon key press, and the subsequent trial began after a randomized inter-trial interval of 4–8 s. This condition served as a control where neither spontaneity nor predictability of action timing was present.

##### Delayed condition (predictable key press)

Each trial commenced when a black dot appeared at a random position on the clock face and began rotating. Participants were instructed to remember this initial position and press the key when the dot completed exactly one full rotation and returned to the starting position. The rotating dot disappeared immediately upon key press, and the subsequent trial began after a randomized inter-trial interval of 4–8 s. This condition allowed participants to predict the precise timing of their action while removing the self-initiated decision of when to act.

##### Self-initiated condition (spontaneous key press)

Each trial commenced when a black dot appeared at a random position on the clock face and began rotating. Participants were instructed to allow the dot to complete at least a half rotation before freely pressing the key at their chosen moment. Immediately after the key press, the rotating dot disappeared, and the subsequent trial began after a randomized inter-trial interval of 4–8 s. Consistent with prior research (Libet et al., 1983; Park et al., 2020; Shibata & Ohira, 2025), participants were explicitly instructed to avoid preselecting the dot’s location for pressing the key, refrain from maintaining identical intervals between trials, and respond spontaneously.

The three conditions were applied in a block design with counterbalanced order across participants. Each condition comprised 50 experimental trials, preceded by practice trials. Participants were offered rest periods between blocks. The timings of dot appearance and key press were recorded and transmitted to the recording software as trigger signals.

#### 2.3.2 Respiration-action naturalness judgment task

Following the main experimental blocks, participants completed an additional task to assess their subjective experience of the relationship between respiration and voluntary movement. Participants were instructed to press the key at their own pace while inhaling for five trials and while exhaling for five trials (order counterbalanced across participants). After completing each set, participants rated the naturalness of pressing the key during each respiratory phase using visual analog scales (VAS). Additionally, participants indicated their overall preference between inhaling and exhaling during key press on a separate VAS (0 = easier to press while inhaling, 100 = easier to press while exhaling).

#### 2.3.3 Questionnaires

Participants completed a series of questionnaires to assess interoceptive awareness, body awareness, and trait anxiety. Interoceptive awareness was measured using the Japanese version of the Multidimensional Assessment of Interoceptive Awareness (MAIA) (Mehling et al., 2012; Shoji et al., 2018). Body awareness was assessed using the Japanese version of the Body Perception Questionnaire-Body Awareness Very Short Form (BPQ-BA-VSF) (Kobayashi et al., 2021; Porges, 1993). Trait anxiety was measured using the Japanese version of the State-Trait Anxiety Inventory (STAI) (Shimizu & Imaei, 1981; Spielberger et al., 1971). Additionally, activity frequency ratings (sports, music, exercise) and the Ten-Item Personality Inventory (Gosling et al., 2003; Oshio & Abe, 2012) were collected for exploratory purposes but not analyzed in the current study.

### 2.4 Procedure

Prior to the experimental session, participants completed a series of questionnaires online. Upon arrival at the laboratory, participants were seated comfortably in front of a computer monitor (EV2495-BK, EIZO) positioned 57 cm away. Participants were informed that the study investigated the timing of voluntary actions, but were not informed about the specific hypothesis regarding respiration-action coupling to avoid any conscious alteration of their breathing patterns. After setting up the physiological recording equipment (respiration belt and ECG electrodes), participants completed the Libet clock task under three conditions (Self-initiated, Delayed, and Immediate), followed by the Respiration-action naturalness judgment task. Stimulus presentation and response collection were controlled using PsychoPy (version 2022.10). The session also included an additional pre-registered task (AsPredicted #240674) not reported in the present study. After completing all tasks, participants were asked whether they had used the timing of their respiration or heartbeat when pressing the key in the Libet clock task. Finally, participants received a debriefing regarding the study’s aims and procedures. The total experimental session lasted approximately 90 minutes.

### 2.5 Data processing

#### 2.5.1 Exclusion criteria for behavioral data

In the Libet clock task, trials were excluded based on condition-specific criteria. In the Self-initiated condition, trials were excluded if participants responded before the rotating dot completed half a rotation (< 2,560 ms from dot onset). In the Delayed condition, trials were excluded if the key press deviated by more than 1,000 ms from the target timing (i.e., one full rotation after dot onset). In the Immediate condition, trials were excluded if the reaction time exceeded 1,000 ms from dot onset. Participants with fewer than 20 valid trials in a given condition were excluded from the circular analysis for that condition. For analyses requiring all three conditions (e.g., condition comparisons), participants with <20 valid trials in any condition were excluded.

#### 2.5.2 Pre-processing for physiological data

Respiratory data were band-pass filtered between 0.2 and 0.8 Hz (Park et al., 2020, 2022). To determine instantaneous respiratory phase, the Hilbert transform was applied to the filtered respiratory signal, yielding phase angles ranging from −π to π. Peaks were defined as phase 0 and troughs as phase ±π; accordingly, inhalation was classified as the trough-to-peak transition (phase −π to 0) and exhalation as the peak-to-trough transition (phase 0 to π). Respiratory data were further cleaned using the following procedures. First, respiratory pause periods, defined as segments where the smoothed respiratory signal (450-ms moving average) remained relatively flat (absolute slope < 0.2 SD of the slope distribution) for at least 500 ms, were identified and excluded from analysis (Harting et al., 2025). Second, respiratory cycles with abnormal durations were excluded based on participant-specific criteria: cycles with total duration, inhalation duration, or exhalation duration outside the range of Q1 – 2.5 × IQR to Q3 + 2.5 × IQR were removed. Finally, the respiratory phase at the time of each behavioral event (dot onset and key press) was extracted for subsequent analyses.

ECG signals were band-pass filtered between 0.5 Hz and 40 Hz (Kligfield et al., 2007). R-peaks were detected, and instantaneous cardiac phase was calculated by linearly interpolating between consecutive R-peaks, with each R-peak assigned a phase of 0 and the subsequent R-peak assigned a phase of 2π. Abnormal R-R intervals, defined as intervals falling outside the range of Q1 − 2.0 × IQR to Q3 + 2.0 × IQR, were identified and excluded for each participant. Subsequently, the cardiac phase at the time of each behavioral event (dot onset and key press) was extracted for further analyses. All physiological data were preprocessed using MATLAB (R2025a).

### 2.6 Statistical analysis

#### 2.6.1 Respiration-action coupling analysis

##### Circular uniformity test

We investigated the synchronization between voluntary movements and respiratory phases (Park et al., 2020). The Hodges–Ajne (omnibus) test was used to determine whether the phase distribution of key presses deviated significantly from uniformity (Ajne, 1968). Given that exhalation typically lasts longer than inhalation, random events tend to cluster during exhalation. To control for this bias, we generated 1,000 surrogate datasets by randomly shifting the respiratory phase data for each participant, allowing the empirical determination of chance-level distributions. For each participant, the statistical value M was computed, defined as the minimum number of data points falling within any half-circle. Lower M values indicate greater deviation from uniformity. To obtain a group-level statistic, M values were summed across participants. The significance of the observed summed M was evaluated using a one-tailed permutation test, as lower M values indicate greater deviation from uniformity. The p-value was computed as the proportion of surrogate summed M values less than or equal to the observed value. All p-values were corrected for multiple comparisons using the false discovery rate (FDR) method. Results for dot onset are reported in the Supplementary Materials (S2).

##### Pairwise Phase Consistency (PPC) analysis

To examine the time course of respiration-action coupling, we calculated Pairwise Phase Consistency (PPC) as a measure of phase concentration that is unbiased by trial number (Vinck et al., 2010). PPC values were computed at each time point within a window of −1 to +1 s around key press at a sampling rate of 200 Hz. This 2-second epoch allows for the characterization of respiration-action coupling dynamics extending from the pre-movement preparatory phase to the post-movement period. Baseline PPC was estimated for each participant by randomly sampling N phase values (matched to the number of valid trials) from the task period, excluding ±250 ms around each event. This procedure was repeated 1,000 times, and the mean was used as the baseline estimate. Baseline-corrected PPC (ΔPPC) was calculated by subtracting baseline PPC from the raw PPC time series. To assess whether ΔPPC significantly exceeded baseline over time in each condition, we employed cluster-based permutation tests (Maris & Oostenveld, 2007), which control for multiple comparisons across time points. Time points where the *t*-statistic (one-sample *t*-test against zero) exceeded the threshold corresponding to p < .05 (one-tailed, testing for ΔPPC > 0) were clustered based on temporal adjacency, and cluster-level statistics were computed by summing the t-values within each cluster. The significance of each cluster was determined by comparing the observed cluster statistic against a null distribution generated from 5,000 sign-flip permutations. Results for dot onset are reported in the Supplementary Materials (S2).

##### Exhalation ratio analysis

Respiratory phases were classified into inhalation (phase −π to 0) and exhalation (phase 0 to π). For each event (dot onset and key press), we calculated the proportion of events occurring during exhalation. Because the inhalation ratio is directly complementary to the exhalation ratio (i.e., inhalation ratio = 1 − exhalation ratio), only the exhalation ratio was used in subsequent analyses. To account for individual differences in baseline exhalation duration, we computed chance-corrected exhalation ratios by subtracting the baseline proportion of time spent in exhalation from the observed exhalation ratio. A 2 (event: dot onset, key press) × 3 (condition: Immediate, Delayed, Self-initiated) repeated-measures ANOVA was conducted on the chance-corrected exhalation ratios. Post-hoc pairwise comparisons between conditions were performed using paired t-tests, and one-sample t-tests comparing each condition to baseline (zero) were performed separately for key press timing. All p-values were corrected using the Holm method.

#### 2.6.2 Respiratory changes associated with key press

To examine whether key press was accompanied by changes in breathing patterns, we analyzed respiratory interval (cycle duration) and respiratory depth (amplitude) around the time of key press.

##### Respiratory interval analysis

For each trial, we extracted three consecutive respiratory cycles: the cycle immediately preceding key press (pre), the cycle during which key press occurred (act), and the cycle immediately following key press (post). A linear mixed-effects model was conducted using the lme4 package (Bates et al., 2015) with condition (Immediate, Delayed, Self-initiated), timing (pre, act, post), and respiratory phase at key press (inhalation, exhalation) as fixed effects, and participant as a random intercept. Degrees of freedom and p-values for F-tests were estimated using the Satterthwaite approximation (Kuznetsova et al., 2017). To decompose significant interactions, we computed difference-in-differences contrasts examining whether the pre-to-post change differed between inhalation and exhalation trials for each condition. Contrasts were tested using asymptotic z-ratios with the emmeans package (Lenth, 2023). Post-hoc comparisons were corrected using the Holm method.

##### Respiratory depth analysis

Respiratory depth was quantified as the z-scored peak-to-trough amplitude of each half-cycle (inhalation or exhalation). For each trial, we extracted three consecutive half-cycles around key press (pre, act, post). The same linear mixed-effects model and contrast analysis were applied as for respiratory interval.

#### 2.6.3 Individual differences analysis

##### Subjective naturalness of respiration-action coupling

To examine whether participants subjectively experienced a preference for pressing the key during a particular respiratory phase, we analyzed responses from the respiration-action naturalness judgment task. First, because the preference ratings deviated from normality (Shapiro-Wilk test, *p* = .002), a Wilcoxon signed-rank test was conducted to determine whether the preference rating (0 = easier to press while inhaling, 100 = easier to press while exhaling) differed significantly from the midpoint (50). Second, we calculated the correlation between the subjective naturalness rating of pressing during exhalation and the exhalation bias (chance-corrected exhalation ratio at key press) in the Self-initiated condition, to examine whether implicit respiratory-action coupling was related to explicit subjective experience.

##### Predictors of coupling strength

To examine individual difference factors associated with respiration-action coupling, we conducted a multiple regression analysis with coupling strength in the Self-initiated condition as the dependent variable. Coupling strength was operationalized as baseline-corrected PPC (ΔPPC) at key press. Interoceptive awareness (MAIA mean score), body awareness (BPQ-BA-VSF sum score), and trait anxiety (STAI-T sum score) were entered as predictors. All variables were standardized prior to analysis.

#### 2.6.4 Exploratory analysis

To examine whether key press timing was coupled with cardiac phase, we applied the same circular analyses used for respiration (Hodges-Ajne test with permutation testing and FDR correction) to cardiac phase data.

To examine whether the mean respiratory phase at key press shifted from baseline, we applied a randomized version of Moore’s paired circular test (Grund et al., 2022; Moore, 1980). For each participant, baseline mean phase was estimated by randomly sampling N phase values (matched to the number of valid trials) from the task period, excluding ±250 ms around each event. This procedure was repeated 1,000 times, and the circular mean of the sampled phases was averaged across iterations to obtain the baseline estimate. Moore’s test compared the observed mean phase at key press against this baseline for each condition, using 10,000 random permutations. All resulting p-values were corrected using the Holm method.

Exploratory analyses examining whether physiological states (respiratory phase, cardiac phase, respiratory interval, and R-R interval) predicted response timing are reported in Supplementary Materials (S3).

Circular statistical analyses were performed using MATLAB (R2025a) and the Circular Statistics Toolbox (Berens, 2009). All other statistical analyses were conducted using R (version 4.3.1).

## 3 Results

### 3.1 Basic measurements

As noted in Section 2.1, one participant was excluded from the Delayed condition due to fewer than 20 valid trials remaining after exclusion criteria were applied; analyses involving all three conditions therefore included 28 participants. Across all participants, the mean exclusion rates were 12.1% (*SD* = 10.8%) in the Self-initiated condition, 15.8% (*SD* = 19.7%) in the Delayed condition, and 9.0% (*SD* = 6.8%) in the Immediate condition. The final mean numbers of valid trials per participant were 44.0 in the Self-initiated condition, 42.1 in the Delayed condition, and 45.5 in the Immediate condition. For respiratory data, approximately 7.7% (*SD* = 2.2%) of respiratory cycles were excluded based on the IQR criterion, and an additional 6.8% (*SD* = 2.8%) were excluded due to respiratory pause periods. For cardiac data, 2.4% (*SD* = 2.8%) of R-R intervals were excluded as outliers.

The mean reaction time in the Immediate condition was 0.342 s (*SD* = 0.067). The mean interval from dot onset to key press was 5.124 s (*SD* = 0.173) in the Delayed condition and 5.808 s (*SD* = 3.053) in the Self-initiated condition. The mean respiratory intervals were 3.36 s (*SD* = 0.57) in the Self-initiated condition, 3.29 s (*SD* = 0.55) in the Delayed condition, and 3.25 s (*SD* = 0.59) in the Immediate condition.

### 3.2 Respiration-action coupling

We tested whether key press timing systematically aligned with spontaneous respiratory phases across the three conditions (Figure 2). In the Self-initiated condition, significant respiratory synchronization was observed at key press (*p* = .009, FDR-corrected), indicating that voluntary key presses were not uniformly distributed across the respiratory cycle. By contrast, no significant synchronization was found in the Delayed condition (*p* = .284) or the Immediate condition (*p* = .166). Results for dot onset timing are reported in Supplementary Materials (S2).

**Figure 1.**
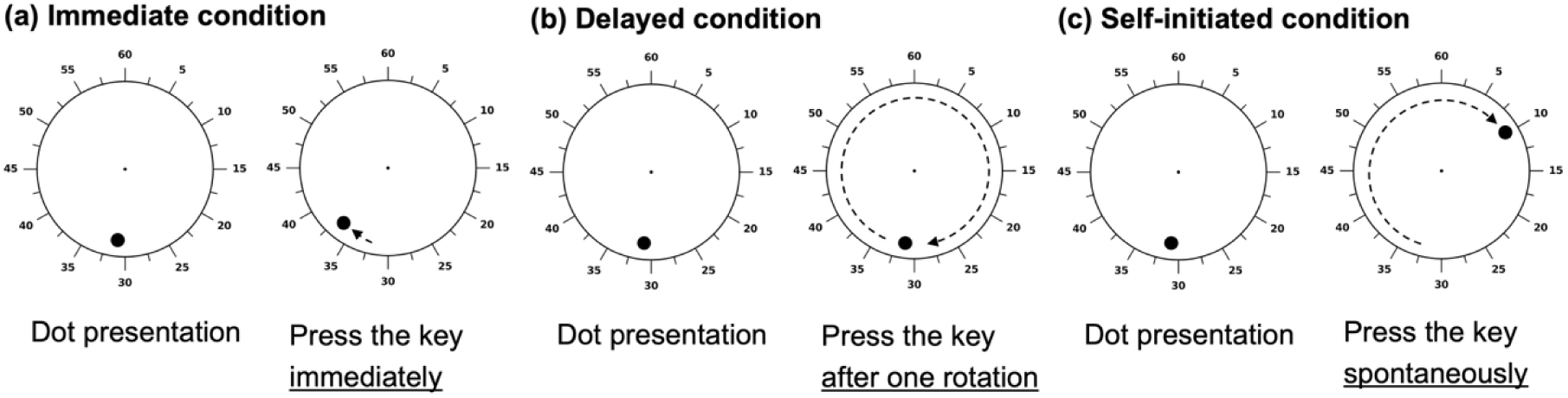
Schematic illustration of the three experimental conditions in the Libet clock task. A dot rotated around a clock face at a constant speed (5,120 ms per rotation). (a) Immediate condition: participants pressed the key immediately upon dot appearance. (b) Delayed condition: participants pressed the key when the dot returned to its initial position after exactly one full rotation. (c) Self-initiated condition: participants pressed the key spontaneously at a self-chosen moment after the dot completed at least half a rotation. Dashed arrows indicate the trajectory of the rotating dot from presentation to key press.

**Figure 2.**
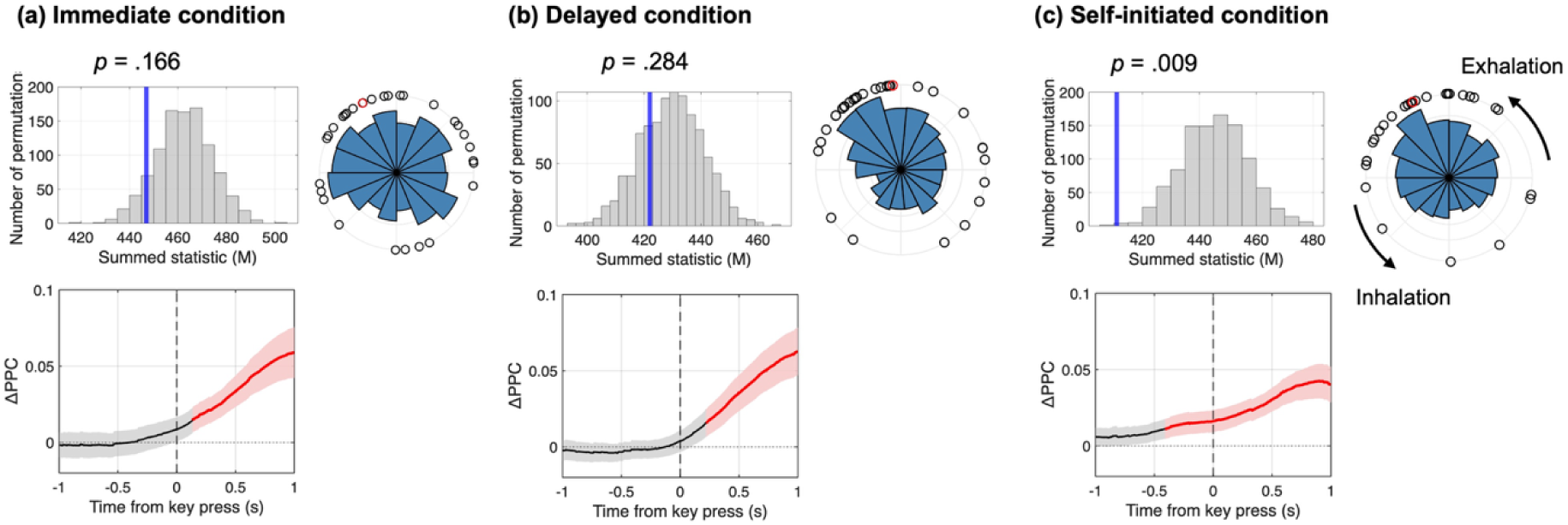
Respiration-action coupling across the three experimental conditions. (a) Immediate condition, (b) Delayed condition, (c) Self-initiated condition. Upper panels: Results of permutation tests for circular uniformity (left) and circular histograms of respiratory phase at key press (right). In the permutation histograms, grey bars represent the null distribution of the summed M statistic, and blue vertical lines indicate the observed values. In the circular histograms, blue bars show the distribution of respiratory phases across all trials. The black dots represent individual participants’ mean respiratory phases, and the red dots indicate the overall group-level mean phase. The arrow in (c) indicates the direction of the respiratory cycle from inhalation to exhalation. Lower panels: Time course of pairwise phase consistency (ΔPPC) relative to baseline around key press. Red segments indicate time periods with significantly elevated PPC identified by cluster-based permutation tests (*p* < .05), and black segments indicate non-significant periods. Shaded areas represent 95% confidence intervals. Only the Self-initiated condition showed significant respiratory-action coupling both in the circular uniformity test and before key press onset in the PPC analysis.

The PPC time-series analysis revealed distinct temporal patterns of respiration-action coupling across conditions (Figure 2). Cluster-based permutation tests identified significant clusters where PPC values exceeded baseline in all three conditions: Self-initiated (−0.41 to 1.00 s, *p* = .004), Delayed (0.22 to 1.00 s, *p* = .024), and Immediate (0.14 to 1.00 s, *p* = .014). Critically, in the Self-initiated condition, the significant cluster began before key press onset, suggesting that respiratory-action coupling emerged before the motor response. In contrast, in the Delayed and Immediate conditions, significant clusters emerged only after key press, likely reflecting respiratory changes induced by the motor action itself.

A 2 (event: dot onset, key press) × 3 (condition: Self-initiated, Delayed, Immediate) repeated-measures ANOVA on chance-corrected exhalation ratios revealed a significant main effect of condition (*F*(2, 54) = 4.43, *p* = .017, 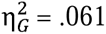), with no significant main effect of event (*F*(1, 27) = 0.15, *p* = .706) or interaction (*F*(2, 54) = 0.24, *p* = .785). As the interaction was not significant, post-hoc pairwise comparisons were conducted on exhalation ratios averaged across events. The Self-initiated condition showed a higher exhalation ratio than the Immediate condition (*t*(27) = 2.63, *p*_adj = .042, *d* = 0.50), while other comparisons were not significant (*p*s > .20). Follow-up one-sample t-tests for key press timing showed that the Self-initiated condition did not significantly differ from baseline after correction (*t*(27) = 2.34, *p*_adj = .080, *d* = 0.44), nor did the Delayed (*t*(27) = 1.23, *p*_adj = .460, *d* = 0.23) or Immediate (*t*(27) = −0.61, *p*_adj = .550, *d* = −0.11) conditions (Figure 3). Although the effect size in the Self-initiated condition suggests a tendency toward exhalation-biased responses, substantial individual differences were observed. This variability was further examined in relation to subjective ratings of movement naturalness.

**Figure 3.**
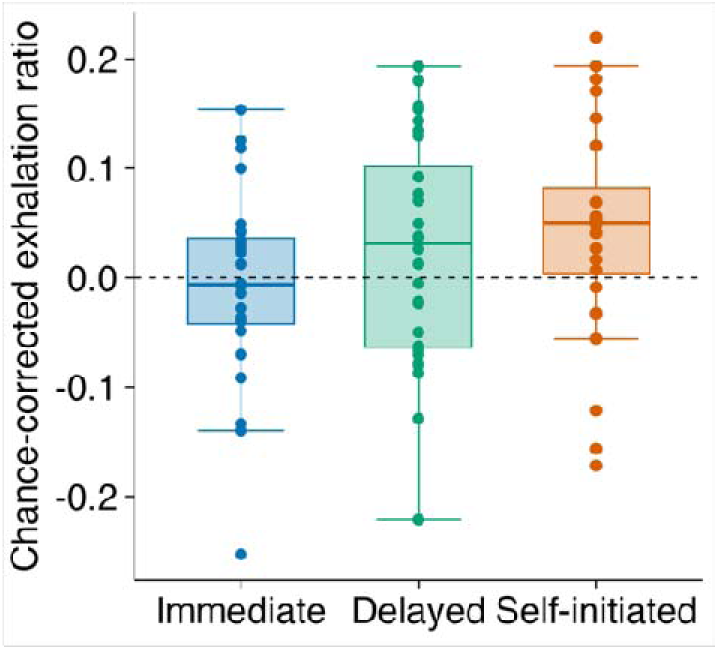
Chance-corrected exhalation ratios at key press across conditions. Box plots show the distribution of chance-corrected exhalation ratios at key press for each condition. Individual dots represent each participant’s data. The dashed line indicates baseline (chance level = 0). Although the Self-initiated condition showed a tendency toward exhalation-biased responses, substantial individual variability was observed.

Together, these results indicate that respiration-action coupling was specific to self-initiated actions. Circular uniformity tests revealed significant phase concentration exclusively in the Self-initiated condition. Crucially, PPC analysis showed that only the Self-initiated condition exhibited a significant cluster before key press onset, suggesting a pre-motor emergence of respiration-action coupling during endogenously timed actions. The directional preference toward exhalation showed substantial individual variability.

### 3.3 Respiratory changes around key press

To examine whether key press was accompanied by changes in breathing patterns, we analyzed respiratory intervals and depths around key press using a linear mixed-effects model (Figure 4).

**Figure 4.**
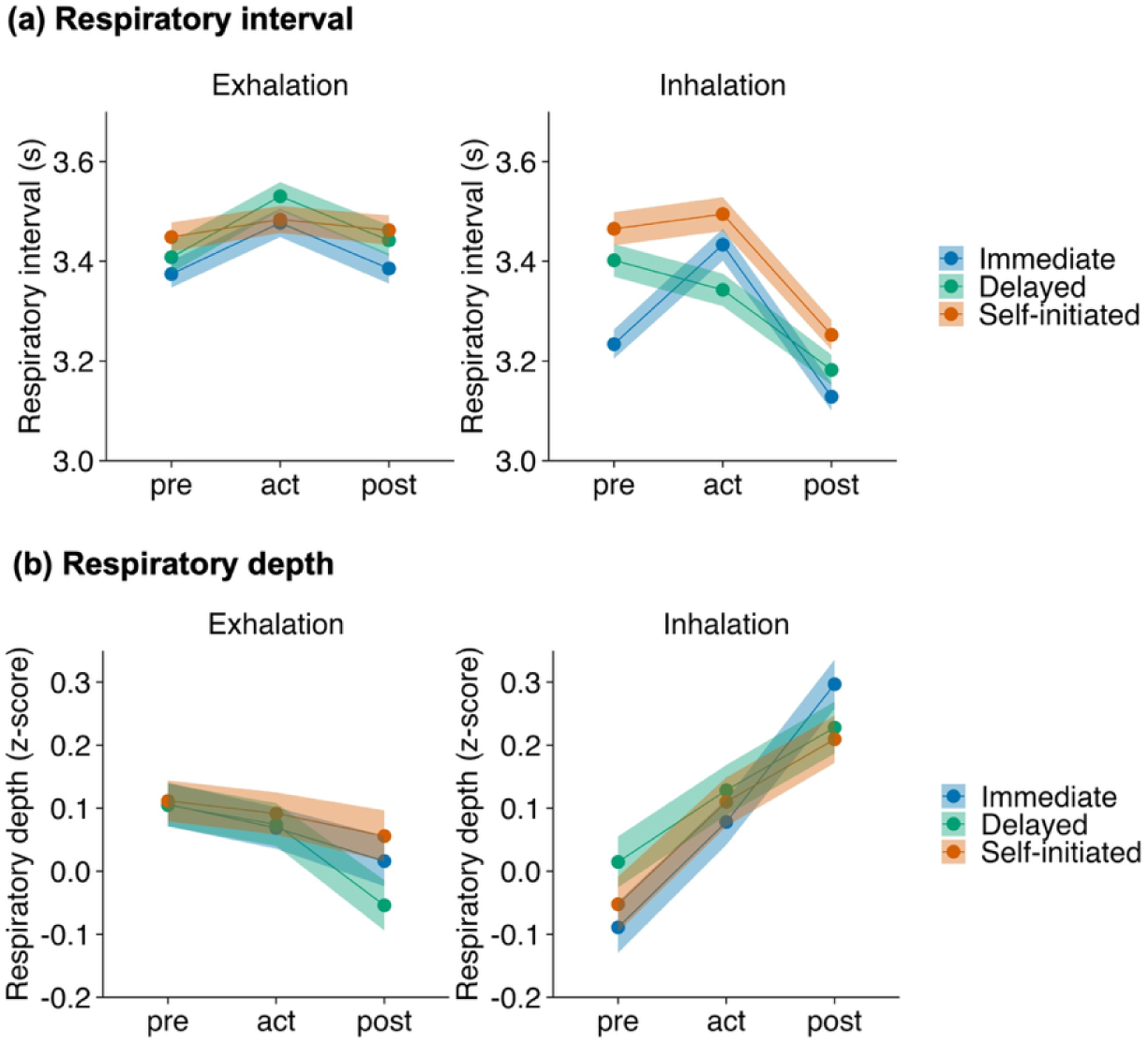
Respiratory changes around key press. (a) Respiratory interval and (b) respiratory depth as a function of timing relative to key press (pre, act, post), separately for trials in which key press occurred during exhalation (left) and inhalation (right). Lines represent mean values for each condition, and shaded areas indicate 95% confidence intervals. When key press occurred during inhalation, breathing became faster (shorter intervals) and deeper in the subsequent cycle, whereas these changes were attenuated when key press occurred during exhalation.

The three-way interaction of condition × respiratory phase × timing was significant (*F*(4, 12356) = 5.47, *p* < .001). To decompose this interaction, we examined whether the pre-to-post change in respiratory interval differed between exhalation and inhalation trials for each condition. This difference was significant in all conditions (Self-initiated: *z* = 4.91, *p* < .001; Delayed: *z* = 5.83, *p* < .001; Immediate: z = 2.65, *p* = .033, Holm-corrected). When key press occurred during inhalation, respiratory intervals shortened after the action, whereas this change was attenuated when key press occurred during exhalation.

We also analyzed respiratory depth (z-scored amplitude) around key press. The respiratory phase × timing interaction was significant (*F*(2, 11484) = 39.53, *p* < .001), while the three-way interaction was not (*F*(4, 11484) = 0.71, *p* = .585), indicating that the relationship between respiratory phase and timing was consistent across conditions. The pre-to-post change in respiratory depth differed significantly between exhalation and inhalation trials in all conditions (Immediate: *z* = 6.36, *p* < .001; Delayed: *z* = 4.90, *p* < .001; Self-initiated: *z* = 4.15, *p* < .001, Holm-corrected). When key press occurred during inhalation, breathing became deeper after the action, whereas this change was attenuated when key press occurred during exhalation.

### 3.4 Individual differences

To examine subjective experience of respiration-action coupling, we first analyzed preference ratings for pressing during exhalation versus inhalation. A Wilcoxon signed-rank test indicated that the preference rating (*Mdn* = 75) was significantly greater than the midpoint of 50 (*V* = 348, *p* = .005, *r* = .52), suggesting that participants generally found it easier to press the key during exhalation (Figure 5a). Furthermore, the subjective naturalness of pressing during exhalation was positively correlated with exhalation bias in the Self-initiated condition (*r* = .46, *p* = .013; Figure 5b), indicating that participants who showed stronger exhalation-biased responses also rated pressing during exhalation as more natural.

**Figure 5.**
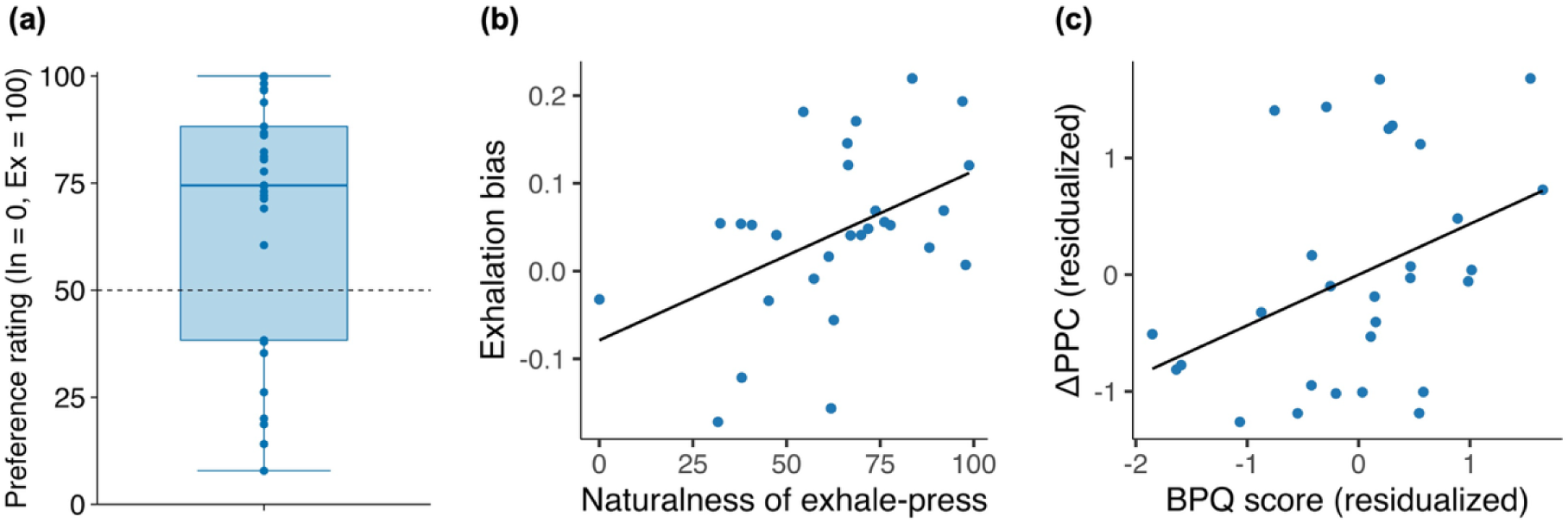
Individual differences in respiration-action coupling. (a) Preference ratings for the ease of pressing during exhalation versus inhalation. Ratings were made on a visual analog scale (0 = easier during inhalation, 100 = easier during exhalation). The dashed line indicates the midpoint (50). Participants generally found it easier to press during exhalation (*Mdn* = 75, *V* = 348, *p* = .005). (b) Correlation between subjective naturalness of pressing during exhalation and exhalation bias in the Self-initiated condition (*r* = .46, *p* = .013). (c) Partial regression plot showing the relationship between body perception (BPQ) and coupling strength (ΔPPC) in the Self-initiated condition, controlling for MAIA and STAI-T (β = .44, *p* = .039). In (b) and (c), each dot represents one participant.

To examine individual difference factors associated with respiration-action coupling, we conducted a multiple regression analysis with coupling strength (ΔPPC) in the Self-initiated condition as the dependent variable, and interoceptive awareness (MAIA), body perception (BPQ), and trait anxiety (STAI-Trait) as predictors. Although the overall model did not reach statistical significance (*F*(3, 25) = 2.60, *p* = .074, *R*² = .24, adjusted *R*² = .15), body perception was a significant predictor of coupling strength (β = 0.44, *t* = 2.18, *p* = .039), indicating that individuals with higher body perception showed stronger respiration-action coupling (Figure 5c). Interoceptive awareness (β = −0.03, *p* = .891) and trait anxiety (β = 0.21, *p* = .270) were not significant predictors.

Finally, to assess whether the observed coupling reflected deliberate strategy, we examined post-experimental reports. Most participants (22/29) reported that they did not consciously use respiratory timing when pressing the key. Seven participants reported some awareness of a relationship between respiration and key press timing; however, most of these reports reflected retrospective awareness (e.g., “it might have happened”) rather than deliberate strategic use. Only two participants explicitly reported intentional use of respiratory timing. Importantly, excluding these two participants did not change the pattern of results (Supplementary Materials; S4), suggesting that the observed coupling was predominantly implicit.

### 3.5 Exploratory analyses

To examine whether key press timing was coupled with cardiac phase, we applied the same circular analyses used for respiration. No significant coupling between cardiac phase and key press timing was observed in any condition (all *p*s > .22, FDR-corrected).

Moore’s paired tests with Holm correction revealed that the mean respiratory phase at key press differed significantly from baseline in the Delayed condition (*R* = 0.46, *p* < .001) and Immediate condition (*R* = 0.42, *p* < .001), but not in the Self-initiated condition (*R* = 0.55, *p* = .098). In the Delayed and Immediate conditions, key presses occurred earlier in the respiratory cycle (toward the inhalation-to-exhalation transition) compared to baseline.

## 4 Discussion

This study investigated the coupling between respiration and voluntary action, exploring its underlying mechanisms and individual differences. We found that (i) respiration–action coupling emerged before action onset exclusively in the Self-initiated condition, (ii) actions performed during inhalation were followed by changes in subsequent respiratory depth and rhythm, and (iii) the direction of synchronization was associated with subjective preference, while coupling strength correlated with self-reported body awareness.

### 4.1 Does action adapt to breathing, or breathing to action?

The present findings suggest that pre-movement respiration–action coupling reflects voluntary action timing aligning with natural breathing rhythms, rather than breathing adjusting to action. Replicating previous findings (Park et al., 2020, 2022; Shibata & Ohira, 2025), respiration–action coupling was observed in the Self-initiated condition. Critically, we extended these findings by introducing a Delayed condition, in which action timing was predictable but not freely chosen, providing sufficient preparation time while removing the freedom to choose when to act. In the Delayed condition, because stimulus onset occurred at random respiratory phases and action timing was fixed relative to the stimulus, coupling could only emerge through respiratory adjustment. However, no such coupling at key press was observed in the Delayed or Immediate condition. Time-series analysis of phase consistency further supported this interpretation: only the Self-initiated condition showed coupling emerging before movement onset, whereas increases in other conditions appeared only after action, likely reflecting motor-induced respiratory changes. These results indicate that predictability alone is insufficient—temporal freedom to choose when to act is necessary for respiration–action coupling to emerge before movement onset. Furthermore, excluding participants who reported intentional breathing control did not alter this pattern, suggesting that coupling operates implicitly.

Previous studies on respiration–action coupling fall into two categories: self-paced tasks where participants freely choose when to act (Park et al., 2020, 2022; Perl et al., 2019; Rassler, 2000), and externally-triggered tasks where actions are cued by stimuli (Chalas et al., 2025; Grund et al., 2022; Harting et al., 2025; Johannknecht & Kayser, 2022; Saltafossi et al., 2025). In self-paced tasks, it has been difficult to determine whether coupling reflects action timing adapting to breathing or breathing adapting to anticipated action, as bidirectional interactions have been suggested (Rassler, 2000; Rassler & Raabe, 2003). Our study directly compared self-chosen timing (Self-initiated) with fixed timing (Delayed and Immediate). Although the Delayed condition provided preparation time and predictability comparable to the Self-initiated condition, pre-movement coupling emerged only when participants could freely choose when to act. This indicates that self-chosen timing, rather than preparation time or predictability, is the critical factor for respiration–action coupling. These findings highlight the importance of temporal freedom in respiration–action coupling (Galvez-Pol et al., 2025; Lopez-Martin et al., 2026), as self-chosen timing provides the flexibility to select action onset based on ongoing internal states, exploiting natural physiological fluctuations.

When action timing is externally fixed, coupling can only emerge through respiratory adjustment. However, such adjustment does not always occur—raising the question of when and why breathing adapts to external events. Functionally, respiratory adjustments can be broadly categorized into two types (Kayser et al., 2025) : passive entrainment and active alignment. Passive entrainment refers to the phenomenon whereby breathing spontaneously synchronizes with regular stimuli or rhythmic movements, resembling a physical coupling process (Goheen et al., 2024; Rassler & Raabe, 2003; Tort et al., 2025). Active alignment, by contrast, involves deliberate respiratory adjustment driven by task demands and performance optimization. In externally-triggered tasks with high performance demands (e.g., detection thresholds, speeded responses), participants may be motivated to adjust their breathing to optimize performance, given that task performance varies across the respiratory cycle (Kluger et al., 2021; Perl et al., 2019; Zelano et al., 2016). At the same time, altering breathing patterns incurs a metabolic cost, as breathing requires continuous muscular work and is homeostatically regulated (Del Negro et al., 2018). Whether respiratory adjustment occurs may therefore depend on the balance between this physiological cost and the potential benefit for task performance. In our Delayed condition, although temporal precision was required, this demand was likely insufficient to motivate the cost of altering natural breathing patterns, which may explain why respiratory adjustment was not observed.

### 4.2 Respiratory changes following action depend on breathing phase: An active inference perspective

Our results revealed that key presses occurring during inhalation were followed by shortened and deepened subsequent breaths, whereas this pattern was attenuated when key presses occurred during exhalation. This qualitative pattern was observed across all three conditions, suggesting that it reflects a fundamental respiratory–motor interaction rather than a task-specific effect. From a physiological perspective, inhalation is an active phase driven by the contraction of inspiratory pump muscles, particularly the diaphragm, and involves considerable metabolic cost (Del Negro et al., 2018). By contrast, exhalation is largely passive and energetically inexpensive, relying primarily on elastic recoil. We interpret the observed post-action respiratory changes as compensatory responses to actions executed during active inhalation.

These findings are consistent with both the motor competition hypothesis and the active inference framework. Previous studies have shown that voluntary key presses tend to be coupled with the exhalation phase (Park et al., 2020, 2022; Shibata & Ohira, 2025); a similar tendency was also observed in the present study. The motor competition hypothesis explains this pattern by positing that actions occur during exhalation to avoid interference between skeletal motor activity and respiratory movements (Park et al., 2020, 2022; Sandoval et al., 2025). From a neural perspective, these studies have also linked the readiness potential to the exhalation phase, although this relationship remains debated (Jeay-Bizot et al., 2026). However, the functional significance of this exhalation preference—why acting during exhalation might be beneficial—has remained unclear. The present results are consistent with this possibility: acting during inhalation was followed by larger respiratory perturbations, suggesting greater interoceptive cost. Of note, the present biphasic analysis cannot distinguish between “selecting exhalation” and “avoiding late inhalation“; the latter may be more accurate, given that late inhalation likely represents a period of maximal respiratory afferent input. The active inference framework offers a complementary perspective: organisms act to minimize prediction error, including interoceptive prediction error (Allen et al., 2023; Barrett & Simmons, 2015; Corcoran et al., 2018). In self-initiated actions, where no external constraints determine action timing, individuals may implicitly select moments that minimize the impact on ongoing breathing. These findings suggest a potential mechanism that may contribute to the exhalation bias observed in voluntary action.

### 4.3 Individual differences in respiration–action coupling: Subjective experience and body awareness

Consistent with previous findings (Shibata & Ohira, 2025), we observed substantial individual variability in respiration–action coupling, both in its direction (preference for inhalation vs. exhalation) and strength. The present study examined whether this variability relates to subjective experience and body awareness. First, in the naturalness judgment task, participants reported a significant preference for pressing the key during exhalation rather than inhalation. Furthermore, subjective naturalness of action during exhalation was positively correlated with the implicit exhalation bias observed in the Self-initiated condition. This suggests a correspondence between explicit preference and implicit coupling, though the direction of this relationship remains to be clarified. Individuals may find exhalation-timed actions more natural because they implicitly tend to act during exhalation, or the subjective preference may drive the behavioral tendency. Second, in the multiple regression analysis, although the overall model did not reach statistical significance, BPQ score significantly predicted the strength of respiration–action coupling in the Self-initiated condition. In contrast, MAIA and STAI scores were not significant predictors. This dissociation may reflect the different aspects of interoception captured by these measures: BPQ assesses the awareness of bodily sensations, whereas MAIA emphasizes metacognitive awareness and regulatory aspects of interoception (Murphy et al., 2020). The tendency to notice bodily sensations, rather than the ability to reflect on them, may be more relevant to respiration–action coupling. Speculatively, individuals with higher body awareness may be more attuned to respiratory signals due to higher precision weighting (Ainley et al., 2016), resulting in stronger respiration–action coupling.

### 4.4 Limitations

First, the trial structure—in which the dot began rotating before participants pressed the key—may have influenced respiratory phase at the time of action (Shibata & Ohira, 2025). Although no significant synchronization between dot onset and respiratory phase was observed (Supplementary Materials S2), in the Delayed and Immediate conditions, the mean respiratory phase at key press was significantly shifted from baseline (Moore’s test), potentially reflecting entrainment to the external stimulus structure. By contrast, the Self-initiated condition showed phase concentration without a significant phase shift, suggesting that coupling in this condition reflects internal selection rather than stimulus-driven entrainment. How stimulus structure and self-chosen timing jointly influence respiratory coupling remains to be fully clarified. Second, the biphasic classification (inhalation vs. exhalation) may obscure finer-grained phase effects. In the present study, the exhalation bias did not survive correction for multiple comparisons, despite a medium effect size (*d* = 0.44). Recent evidence suggests that voluntary actions cluster during late exhalation to early inhalation (Jeay-Bizot et al., 2026), indicating that effects may be concentrated at phase transitions rather than within phases per se. Such patterns would not be captured by simple biphasic classification, and future studies with finer-grained phase analyses are warranted. Third, the sample size (*N* = 29) was determined based on phase coupling analyses and may have been underpowered for detecting individual difference effects. Although BPQ significantly predicted coupling strength, the overall regression model did not reach statistical significance, and these findings should be interpreted cautiously pending replication with larger samples. Finally, in the naturalness judgment task, participants were explicitly instructed to press the key while attending to their breathing phase, which may not reflect the implicit respiration–action relationship observed under natural conditions. The correspondence between subjective preference and implicit coupling should therefore be interpreted with this methodological constraint in mind.

## 4.5 Conclusion

In conclusion, the present study provides evidence that pre-movement respiration–action coupling reflects voluntary action timing aligning with natural breathing rhythms, rather than breathing adjusting to action. This coupling emerges implicitly when individuals have temporal freedom, suggesting that self-chosen timing allows for the exploitation of ongoing physiological states. The finding that actions during inhalation were followed by greater respiratory changes offers a potential functional explanation for the widely observed exhalation bias in voluntary action: acting during exhalation may minimize interoceptive cost. Furthermore, the correspondence between implicit coupling and subjective preference, along with the association between coupling strength and body awareness, suggests that respiration–action coupling reflects individual differences in interoceptive processing. These findings contribute to our understanding of how internal physiological states shape the timing of voluntary behavior.

## Author Contributions

**Hiroshi Shibata**: Conceptualization, Methodology, Data curation, Formal analysis, Investigation, Writing – original draft, Writing – review and editing, Visualization. **Hideki Ohira**: Conceptualization, Methodology, Supervision, Writing – review and editing, Funding acquisition.

## Data and Code Availability Statement

The data and code that support the findings of this study will be made publicly available upon publication of this article.

## Declaration of Generative AI and AI-assisted technologies in the writing process

During the preparation of this work the authors used Claude (Claude Opus 4.5, Anthropic) in order to correct English language. After using this tool, the authors reviewed and edited the content as needed and take full responsibility for the content of the published article.

## Acknowledgments

We thank the undergraduate students of our psychology course for their help with data collection.

