## Supplementary Materials for "Pre-movement respiration–action coupling is specific to self-initiated action: Evidence for action alignment with ongoing breathing"

**S1. Deviations from preregistration.**

This study was preregistered on AsPredicted (#240669; https://aspredicted.org/8pw9sq.pdf). The following deviations from the preregistration occurred:

First, in addition to the preregistered Hodges–Ajne permutation tests (with FDR correction) on respiratory phases at dot onset and key press, we conducted time-resolved Pairwise Phase Consistency (ΔPPC) analysis with cluster-based permutation testing to examine the temporal dynamics of respiration-action coupling. Although event-locked respiratory changes were preregistered as exploratory analyses, the specific analytical approach (PPC with cluster permutation) was determined post hoc.

Second, the preregistration specified that participants providing fewer than 20 valid trials in any condition would be excluded from the study. In practice, such participants were excluded from analyses requiring data from all three conditions (e.g., condition comparisons), but were retained for single-condition analyses (e.g., Self-initiated condition only) when applicable. This modification was made to maximize data retention while maintaining analytical validity.

Third, the preregistered individual-differences analysis specified that respiratory phase concentration (circular concentration) would be predicted by MAIA, BPQ, and STAI-T scores. In the present report, we used ΔPPC at key press in the Self-initiated condition as the measure of respiration-action coupling strength. Although both measures index the coupling between respiration and action, the specific operationalization differs from the preregistration.

Fourth, the Ten-Item Personality Inventory (TIPI-J) and activity frequency ratings (sports, music, exercise) were collected as specified in the preregistration but are not analyzed in the current study; these data will be made available in the open dataset.

None of these deviations affect the interpretation of the main findings, and all analysis scripts will be made publicly available upon publication.

**S2. Respiration-action coupling at dot onset**

We examined whether dot onset timing was coupled with respiratory phase using the same analyses applied to key press timing. Permutation tests for circular uniformity revealed no significant synchronization between dot onset and respiratory phase in any condition (all *p*s ≥ .498, FDR-corrected; Figure S1), indicating that dot presentation occurred uniformly across the respiratory cycle regardless of condition.

The PPC time-series analysis for dot onset showed no significant respiration-coupling effects (Figure S1). Cluster-based permutation tests did not identify any significant clusters in the Self-initiated condition (*p* = .061), Delayed condition (no clusters detected), or Immediate condition (*p* = .062). This pattern contrasts with the key press results and suggests that respiration-action coupling was specifically linked to the timing of voluntary action rather than stimulus onset.

Together, these results confirm that respiratory-action coupling at dot onset was minimal, supporting the interpretation that the coupling observed at key press in the Self-initiated condition reflects synchronization specific to voluntary action initiation rather than stimulus processing.

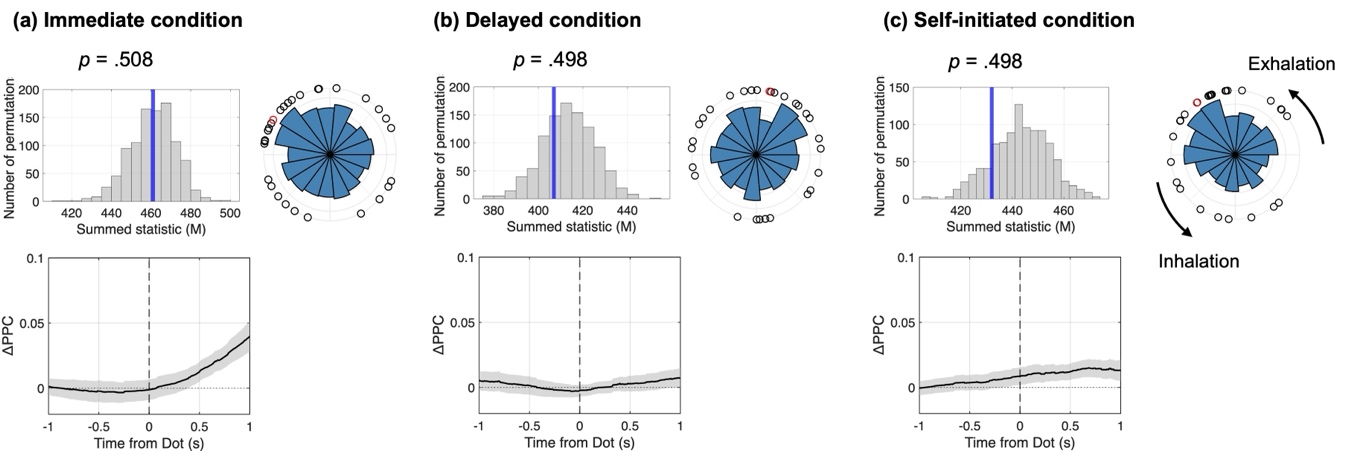

**Figure S1. Respiration-action coupling at dot onset across the three experimental conditions.** Format is the same as Figure 2. No significant respiratory synchronization was observed at dot onset in the circular uniformity tests (all *p*s ≥ .498). Cluster-based permutation tests also did not reveal significant clusters in any condition, indicating that respiration-event coupling was specific to key press timing.

**S3. Physiological predictors of response timing**

To examine whether physiological states predicted response timing, we conducted linear mixed-effects models using the lme4 package (Bates et al., 2015). Reaction time (Immediate condition), interval from dot onset to key press (Self-initiated condition), or timing error from the target timing (Delayed condition) served as the dependent variable. The respiratory interval and R-R interval immediately preceding key press, respiratory phase at key press (cosine and sine components), and cardiac phase at key press (cosine and sine components) were entered as fixed effects, with participant as a random intercept. Separate models were fitted for each condition.

In the Immediate condition, R-R interval significantly predicted reaction time (*β* = −0.096, t = −3.37, *p* < .001), indicating that longer R-R intervals (i.e., slower heart rate) were associated with faster responses. In the Self-initiated condition, R-R interval significantly predicted the interval from dot onset to key press (*β* = 4.03, *t* = 2.84, *p* = .005), with shorter R-R intervals (i.e., faster heart rate) associated with shorter waiting times. The direction of effects thus differed between conditions. In the Delayed condition, none of the predictors significantly predicted timing error (all *p*s > .21). Notably, respiratory interval and respiratory/cardiac phase at key press did not significantly predict response timing in any condition.

**S4. Sensitivity analysis excluding participants who reported intentional use**

Two participants who explicitly reported using respiratory timing were excluded. The pattern of results remained unchanged (Table S1).

**Table S1.** Sensitivity analysis results

| Analysis | Full sample (N = 29) | Excluding intentional users (N = 27) |
| --- | --- | --- |
| Hodges-Ajne (Self-initiated) | *p* = .009 | *p* = .009 |
| Hodges-Ajne (Delayed) | *p* = .284 | *p* = .318 |
| Hodges-Ajne (Immediate) | *p* = .166 | *p* = .211 |
| ΔPPC cluster (Self-initiated) | −0.41 to 1.00 s, *p* = .004 | −0.29 to 1.00 s, *p* = .005 |

*Note.* *p*-values for Hodges-Ajne tests are FDR-corrected. ΔPPC cluster times indicate significant time windows relative to key press.
